# Open Humans: A platform for participant-centered research and personal data exploration

**DOI:** 10.1101/469189

**Authors:** Bastian Greshake Tzovaras, Misha Angrist, Kevin Arvai, Mairi Dulaney, Vero Estrada-Galiñanes, Beau Gunderson, Tim Head, Dana Lewis, Oded Nov, Orit Shaer, Athina Tzovara, Jason Bobe, Mad Price Ball

**Author notes:** Authors contributed equally (Misha Angrist, Kevin Arvai, Mairi Dulaney, Vero Estrada-Galiñanes, Beau Gunderson, Tim Head, Dana Lewis, Oded Nov, Orit Shaer, Athina Tzovara).

## Abstract

**Background:** Many aspects of our lives are now digitized and connected to the internet. As a result, individuals are now creating and collecting more personal data than ever before. This offers an unprecedented chance for human-participant research ranging from the social sciences to precision medicine. With this potential wealth of data come practical problems (such as how to merge data streams from various sources), as well as ethical problems (such as how to best balance risks and benefits when enabling personal data sharing by individuals).

**Results:** To begin to address these problems in real time, we present Open Humans, a community-based platform that enables personal data collections across data streams, giving individuals more personal data access and control of sharing authorizations, and enabling academic research as well as patient-led projects. We showcase data streams that Open Humans combines (e.g. personal genetic data, wearable activity monitors, GPS location records and continuous glucose monitor data), along with use cases of how the data facilitates various projects.

**Conclusions:** Open Humans highlights how a community-centric ecosystem can be used to aggregate personal data from various sources as well as how these data can be used by academic and citizen scientists through practical, iterative approaches to sharing that strive to balance considerations with participant autonomy, inclusion, and privacy.

## Background

Research involving human participants, from biomedical and health research to social sciences studies, is experiencing rapid changes. The rise of electronic records, online platforms, and data from devices contribute to a sense that these collected data can change how research in these fields is performed [1, 2, 3, 4]

Among the impacted disciplines is precision medicine - which takes behavioral, environmental, and genetic factors into account and has become a vision for health-care in the United States [5]. By taking individual parameters into account, precision medicine aims to improve health outcomes, for example by optimizing drugs based on a patient’s genetic makeup [6, 7].

Access to large-scale data sets, along with availability of appropriate methods to analyze these data [8, 9], is often described as a major prerequisite for the success of precision medicine [10]. Falling costs for large-scale, individualized analyses such as whole-genome sequencing [11] have already helped facilitate both research in precision medicine and its adoption. In addition, an increasing number of patients and healthy individuals are collecting health-related data outside traditional healthcare, for example through smartphones and wearable devices [12, 13] or through direct-to-consumer (DTC) genetic testing [14].

Indeed, at least 12-17 million individuals have taken a DTC genetic test [15, 16], while more than 25 million such tests have been purchased [17]. Meanwhile, it is estimated that by 2020 over two exabytes of storage will be needed for health care data [18] alone. Furthermore, data from social network sites like Facebook and Twitter are increasingly likely targets for medical data mining [19]. Additionally, more data is becoming available from personal medical devices, both in real-time and for retrospective analyses [20].

These changes to research and medical practice bring with them a number of challenges, including the problems of data silos, ethical data sharing, and participant involvement. A participant-centered approach to personal data aggregation, sharing, and research has the potential to address these issues. To achieve this, we created *Open Humans* as a digital ecosystem designed to facilitate individual data aggregation across data sources, granular management of data sharing, and co-created research.

### Data Silos

To fully realize the promises of these large personal data collections, not only in precision medicine but all fields of research, access to both big data and smaller data sources is needed, as is the ability to tap into a variety of data streams and link these data [21, 10]. Data silos can hinder the merging and re-use of data by third parties for a number of reasons: they can be incompatible due to different data licenses [22] or inaccessible due to privacy, ethical, and regulatory concerns [23, 24, 25]. For example, the US National Human Genome Research Institute’s Database of Genotypes and Phenotypes remains an underused resource because of logistical and regulatory/ethical oversight challenges for would-be users [26]. In addition to legal barriers, there are typically technical challenges in rendering data accessible, usable, and/or anonymized, and a data controller typically has incentives to seek compensation in return for these activities.

Beyond biomedical datasets, there are data from wearable devices, social media, and other data held by private companies, from which data exports are often not available. In other cases data access might be legally mandated, but the practical outcomes are mixed or in progress [27, 28], *e.g.*, for clinical health data in the United States as mandated by the 1996 *Health Insurance Portability and Accountability Act* and 2009 *Health Information Technology for Economic and Clinical Health Act* (HIPAA and HITECH Act), and for personal data in the European Union as mandated by rights to data access and data portability in the 2016 *General Data Protection Regulation* (GDPR) [29, 30]. In addition, within the context of research involving human participants, data access may be recommended [31] but not legally required, and as a result is not typically provided [32]. Data portability and easy access to research data by participating individuals could empower them to steer research in directions that affect their lives and health outcomes.

### Ethical Data Re-Use

While the sharing and re-using of biomedical data can potentially transform medical care and medical research, it brings along a number of ethical considerations [33, 34]. In the field of human genetics, the ethics of sharing data has been extensively considered with respect to how research participants and patients can give informed consent for studies that carry risks of genetic discrimination, loss of privacy, and re-identification in publicly shared data [35, 36]. Due to access and portability issues, however, research with biomedical data is rarely driven by the individuals from whom the data came – and as a result, such research fails to give patients much power over how their data can be used [37]. For example, it is now abundantly clear that direct-to-consumer genetic testing customers routinely have their de-identified (but re-identifiable) data shared with third parties [38]. Open Humans seeks to be among the agents for change in this regard. Bottom-up research initiatives have included disease- and/or mutation-specific efforts [39, 40] and the development of platforms meant to allow participants to control data-sharing at a granular level [41]. Open Humans is meant to complement such initiatives and enable the creation of multiple “sandboxes” where both personal and biomedical data can be leveraged to help grow empirical knowledge and further downstream development of diagnostics and therapies.

Elsewhere, social media is also gaining importance in research as well as public health [42]. Differing perceptions on the sensitivity of social media data can lead to privacy concerns. For example, an analysis performed on 70,000 users of an online dating website, where private personal data was scraped by researchers and then publicly shared, caused a public outcry [43]. Such cases have sparked calls for caution in performing “big data” research with these new forms of personal data [44, 45].

Research that interacts with social media users raises additional concerns. For example, Facebook was widely criticized for an experiment to study emotional contagion among 700,000 of its users without their consent or debriefing, prompting discussion of the ethics of unregulated human subjects research and “A/B testing” by private entities [46, 47, 48]. And the 2018 disclosure of the Cambridge Analytica controversy, in which a private firm harvested information from 50 million Facebook users without their permission, led Facebook to tighten control over its application programming interfaces (APIs), turning it even more of a silo that does not allow for research to be done by outside researchers [49].

For the foreseeable future, researchers that re-use data from commercial sources will have to decide how to balance the interests of commercial data controllers, participants, and society. While there is no consensus on how research consent for existing personal data should be performed, we know that participants desire more granular abilities to manage data sharing: to decide who can and cannot see it, under what circumstances, and what can and cannot be done with it [50]. Such individual control will be especially critical in the sensitive context of precision medicine [24].

### Participant Involvement

Citizen science mostly describes the involvement of volunteers in the data collection, analysis, and interpretation phases of research projects [51], thus both supporting the research process itself and helping with public engagement. Furthermore, the Universal Declaration of Human Rights describes a broad human right to access science as a whole, implying a right to participate in all aspects of the scientific enterprise [52].

Traditionally, many participatory science projects have focused on the natural sciences, like natural resource management, environmental monitoring/protection, and astrophysics [53, 54, 55]. In many of these examples volunteers are asked to crowd-source and support scientists in the collection of data - e.g. by field observations or through sensors [56] or by performing human computation tasks such as classifying images [57] or generating folded protein-structures [58].

Analogous to the movement in other realms of citizen science, there is a growing movement toward more participant/patient involvement in research on humans, including in fields such as radiology, public health, psychology, and epidemiology [59, 60]. Patients often have a better understanding of their disease and needs than medical/research professionals [61, 62] and that patient involvement can help catalyze policy interventions [63]. Examples include the studies on amyotrophic lateral sclerosis initiated by *PatientsLikeMe* users [64], crowd-sourcing efforts like *American Gut* [65], and a variety of other *citizen genomics* efforts [66]. It is likely that involving patients in clinical research can not only help minimize cost but can lead to drugs being brought to market sooner [67].

Elsewhere, the *Quantified Self* movement, in which individuals perform self-tracking of biological, behavioral, or environmental information and design experiments with an *n=1* to learn about themselves [68], can be seen on this continuum of participant-led research [69]. By performing self-experiments and recording their own data, individuals can gain critical knowledge about themselves and the process of performing research. Analogous to the benefits of patient insights in clinical research, individuals engaged in self-tracking and personal data analysis have the potential to contribute their insights to a variety of other research areas.

### A participant-centered approach to research

As shown above, substantially involving patients and participants in the research process has multiple benefits. Participants as primary data holders can help in breaking down walls among data silos and in aggregating and sharing personal data streams. Furthermore, by being involved in the research process and actively providing data, they can gain autonomy and can actively consent to their data being used, thus mitigating (but not eliminating) the likelihood of subsequent ethical concerns. Last but not least, enabling individuals to analyze and explore their own data, individually and collectively, can result in valuable feedback that helps researchers incorporate the needs, desires, and insights of participants.

In recent years a number of projects have started to explore both data donations and crowd-sourcing research with an extended involvement of participants. In genomics, both academic projects like *DNA.Land* [70] and community-driven projects like *openSNP* [71] are enabling crowdsourcing via personal genetic data set donations. Furthermore, the idea of *Health Data Cooperatives* that are communally run to manage access to health data has emerged [24].

However, most of these projects limit participants’ involvement in the research process: a participant is limited, for example, to providing specific types of data for a specific data repository. Additionally, participants are rarely given an easy way to help in designing a study, let alone running their own.

To close these gaps we developed *Open Humans*, a community-based platform that enables its members to share a growing number of personal data types; participate in research projects and create their own; and facilitate the exploration of personal data by and for the individual member. *Open Humans* was initially conceived as an iteration of work with the Harvard Personal Genome Project [72]. Along with a description of the platform itself and its power and limitations, we present a set of examples on how the platform is already being used for academic and participant-led research projects.

## Results

We designed *Open Humans* as a web platform with the goal of easily enabling connections to existing and newly created data sources and data (re-)using applications. Platform members import data about themselves from various sources into their Open Humans account. They can then explore their aggregated data and share it with projects from citizen scientists and academic researchers.

### Design

In the center of the design are three main components: *Members, Projects* and *Data* objects. *Members* can join various *Projects* and authorize them to read *Data* that’s stored in their account as well as write new *Data* for this *Member* (see Figure 1 for a dataflow diagram).

**Figure 1.**
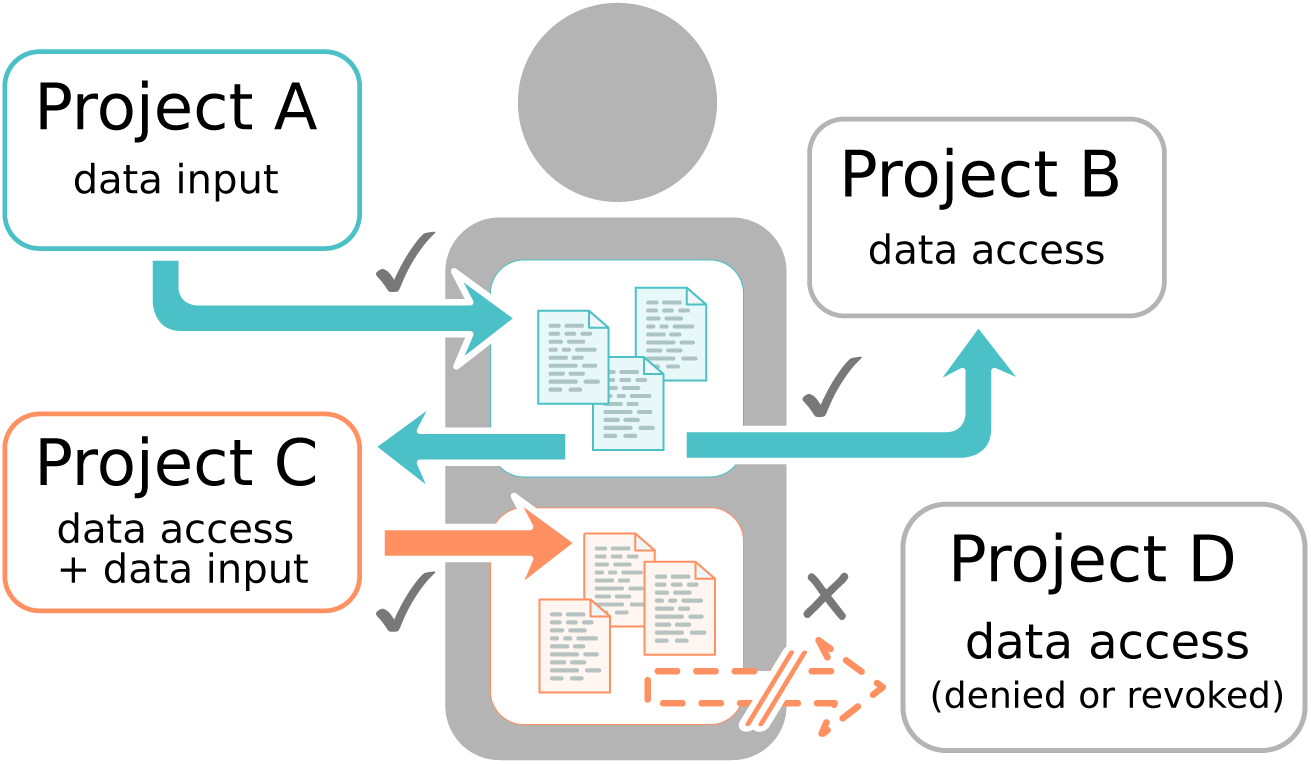
The Open Humans authorization flow. A *Member* (center) can join *Projects* and approve them to read or write *Data*. The *Member* approves *Project A* to deposit files (blue) into their account. They also approve *Project B* to read the files that *Project A* has deposited. Additionally, the *Member* approves *Project C* to both read the files of *Project A* and write new files. The *Member* declines to give access to their personal data to *Project D*.

#### Projects

*Projects* are the primary way for *Members* to interact with *Open Humans*. *Projects* can be created by any *Member*. During project creation a prospective project lead must provide a description of their project and specify the access permissions they request from *Members* who decide to join. These may include:

**Username** By default projects do not get access to a *Member’s* username; each *Member* is identified with a random, unique identifier specific to that project. This way *Member’s* can join a project while being pseudonymous.

**Data Access** A *Project* may ask permission to read *Data* that have been deposited into a *Member’s* account by other projects. A project lead needs to specify to which existing projects’ data they want to have access to and only this data will be shared with the new project.

Through the permission system, *Members* get a clear idea of the amount of *Data* they are sharing by joining a given *Project* and whether their username will be shared and with whom.

Furthermore, all *Projects* have the following permissions for any *Members* who have joined them: (1) they can send messages to *Members*, which are received as emails; and (2) they can upload new *Data* into the *Member* account. Thus, in addition to acting as potential data recipients, *Projects* are also the avenue by which *Data* is added to *Member* accounts.

*Projects* can be set up in two different ways: As an *On-site Project* or as an *OAuth2 Project*. The *OAuth2 Project* format offers a standard OAuth2 user authorization process commonly used to connect across web services. Projects that implement this can connect an Open Humans user to a separate mobile or web application, and can be fully automated. For projects that do not have separate applications, the *On-site Project* format allows the project to present “consent” or “terms of use” information within Open Humans, thereby minimizing the need for technical work on the part of a project. Both formats have access to APIs for performing data uploads, data access, and member messaging.

*Projects* also need to clearly signal whether they are a research study that is subject to ethical oversight by an Institutional Review Board (IRB) or equivalent, or whether they are not performing such research (i.e. not subject to this oversight). This allows for participant-led projects outside an academic research setting, provided *Members* see a notification alerting them to the absence of IRB oversight.

Thus, any Member can create a project in the site, at any time, and all APIs work immediately. However a *Project* will not be publicly listed for *Members* to see, and has a cap limiting the number of *Members* that may join. Public listing and unlimited usage is granted when a project is marked as “approved” following a community review process. *Projects* that have IRB oversight are required to provide documentation of IRB approval as part of this review process.

In summary, given the broad potential features available, a *Project* can cover anything from data import tools, to data processing tools, to research projects, to self-quantification projects that visualize and analyze a *Member’s* data.

#### Members

*Members* interact with *Projects* that are run on *Open Humans*. By joining projects that act as data uploaders, they can add specific *Data* into their Open Humans accounts. This is a way to connect external services: e.g. put their genetic data or activity tracking data into their *Open Humans* account. Once they have connected to relevant *Projects* that import their own data, *Members* can opt-in to joining additional *Projects* that they wish to grant access to their account’s data.

As *Members* are able to selectively join *Projects*, they can elect which projects their *Data* should be shared with. Members may withdraw from a *Project* at any time. This results in immediate revocation of *Data*-sharing authorization for that *Project*, as well as a removal of *Data* upload and message permissions. *Projects* may also support data erasure requests upon withdrawal, and any remaining *Data* uploaded by a project may be retained or deleted by the *Member*. *Open Humans* also allows *Members* to delete their entire account at any time, resulting in an immediate removal from the database, cessation of data processing activities, and permanent deletion following the automated turnover of back-up storage.

#### Data input and management

*Data* is uploaded into a *Member*’s account, which allows any joined *Projects* with requisite permissions to access this data. To be fully available to all of the possible projects that can be run on *Open Humans*, all data are stored in files that can be downloaded by users and *Projects* that have gotten permission. For any file that a *Project* deposits into a *Member*’s account, the uploading *Project* needs to specify at least a description and tags as meta data for the files.

*Members* can always review and access the *Data* stored in their own accounts. By default, the *Data* uploaded into their accounts is not shared with any projects but the one that deposited the data, unless and until other *Projects* are joined and specifically authorized to access this data. In addition to being able to share data with other *Projects*, *Members* can also opt-in into making the data of individual data sources publicly available on a project-by-project basis. *Data* that has been publicly shared is then discoverable through the *Open Humans* Public Data API, and is visible on a *Member*’s user profile.

### Open Humans in Practice

Using this design, the platform now features a number of projects that import data directly into *Open Humans*. Among data sources that can be imported and connected are *23andMe, AncestryDNA, Fitbit, Runkeeper, Withings, uBiome* and a generic *VCF* importer for genetic data like whole exome or genome sequences. Furthermore, as a special category, the *Data Selfie* project allows *Members* to add additional data files that are not supported by a specialized project yet.

The community around the *Open Humans* platform has expanded the support to additional *Data* sources by writing their own data importers and data connections. These include a bridge to *openSNP*, and importers for data from *FamilyTreeDNA, Apple HealthKit, Gencove, Twitter* and the *Nightscout* (open source diabetes) community. Across these data importers, the platform supports data sources covering genetic and activity tracking data as well as recorded GPS tracks, data from glucose monitors, and social media.

The platform has grown significantly since its launch in 2015: As of May 30th 2019, 6,976 members have signed up with *Open Humans*. Of these, 2,945 members have loaded 19,949 data sets into their accounts. In cases where external data sources support the import of historical data (e.g. Fitbit, Twitter), data sets can include data that reaches back before the launch of Open Humans. Furthermore, overall there are now 30 projects that are actively running on Open Humans, with an additional 12 projects that have already finished data collection and thus have been concluded (see Table 1 for the most heavily engaged projects).

**Table 1.** *Open Humans* projects with more than 250 members

| Project name | Description | Members | Data deposited | Data access requested |
| --- | --- | --- | --- | --- |
| 23andMe Upload | Enables members to import their 23andMe data | 1202 | 23andMe data | - |
| Genevieve Genome Report | Matches a member's genome against public variant data, and invites them to contribute to shared notes. | 845 | - | 23andMe Upload, Harvard PGP, Genome/Exome Upload, Username & public data |
| Harvard Personal Genome Project | Enables members to import their data from the Personal Genome Project | 812 | Full genome sequencing data & survey data | - |
| Twitter Archive Analyzer | Enables members to import their Twitter archives and analyzes them | 531 | Twitter archives | - |
| Personal Data Notebooks | Enables personal data analyses with Jupyter Notebooks | 524 | Jupyter Notebooks | All Data |
| Keeping Pace | Seeks to study data about how we move around, to understand how seasons and local environment influence our movement patterns. | 403 | - | Fitbit, Jawbone, Moves, Apple HealthKit, Runkeeper |
| AncestryDNA Upload | Enables members to import their AncestryDNA data | 438 | AncestryDNA data | - |
| Fitbit Connection | Connect a member's Fitbit account to add data from their Fitbit activity trackers and other Fitbit devices. | 404 | Data from a Fitbit account | - |
| GenomiX Genome Exploration | A study of how people interact with their genome data using GenomiX, a visualization tool | 365 | - | Username & public data |
| Circles | A research study that aims to discover the genetic basis for a mysterious and remarkable human trait: the areola. | 321 | - | 23andMe, AncestryDNA, Data Selfies, Harvard PGP, Genome/Exome Upload |
| Gencove | Your genome app - get your ancestry, microbiome, and more! Contribute your data to OpenHumans. | 311 | Sequencing bam files | - |
| openSNP | Enables members to connect their *Open Humans* and *openSNP* accounts | 308 | openSNP user details | Username & public data |
| Nightscout Data Transfer | A tool to easily enable the upload of data from individual Nightscout databases | 293 | Nightscout data | - |

Data was collected on 2019-04-25

### Use Cases

To demonstrate the range of projects made possible through the platform and how the community improves the ecosystem that is growing around *Open Humans*, here we highlight some of the ongoing projects, covering participant-led research, academic research, and projects originating in the self-quantification community.

#### OpenAPS and Nightscout Data & Data Commons

There are a variety of open source diabetes tools and applications that have been created to aid individuals with type 1 diabetes in managing and visualizing their diabetes data from disparate devices. One such tool is Nightscout, which allows users to access continuous glucose monitoring (CGM) data. Another such example is OpenAPS, the Open Source Artificial Pancreas System, which is designed to automatically adjust an insulin pump’s insulin delivery to keep users’ blood glucose in a safe range overnight and between meals [73]. These tools enable real-time and retrospective data analysis of rich and complex diabetes data sets from the real world.

Traditionally, gathering this level of diabetes data would be time-consuming, expensive, and otherwise burdensome to the traditional researcher, and often pose a prohibitive barrier to researchers interested in getting started in the area of diabetes research and development. Using Open Humans, individuals from the diabetes community have created a data uploader tool called Nightscout Data Transfer Tool to enable individuals to share their CGM and related data with the Nightscout and/or OpenAPS Data Commons [74]. Sharing is done pseudonymously via random identifiers, enabling an individual to protect their privacy. Furthermore, sharing is facilitated as a single data upload may be used in multiple studies and projects. These two patient-led data commons have requirements for use that allow both traditional *or* citizen science (e.g. patient) researchers to use this data for research. These data commons were created with the goal of facilitating more access to diabetes data such as CGM datasets that are traditionally expensive to access. By doing so, they enable more researchers to explore innovations for people with diabetes. Additionally, OpenAPS is the first open source artificial pancreas system with hundreds of users, who are hoping such data sharing will facilitate better tools and better innovations for academic and commercial innovations in this space. To date, dozens of researchers and many community members have accessed and utilized data from each of these commons. Some publications and presentations have also showcased the work and the data donated by members of the community, further allowing other researchers to build on this body of work and these data sets [75] (https://openaps.org/outcomes/).

In addition to facilitating easier access to more and richer diabetes data, the Nightscout and OpenAPS communities have also been developing a series of open source tools to enable individuals to more easily work with the datasets (https://github.com/danamlewis/OpenHumansDataTools).

#### Linking across communities: openSNP

*openSNP* is a database for personal genomics data that takes a different approach than *Open Humans*. While *Open Humans* focuses on granular control in terms of whom Members share their data with, *openSNP* focuses on maximizing re-use of data, by exclusively allowing individuals to donate the raw DTC genetic test data into the public domain [71]. With already over 4,500 genetic data sets, *openSNP* is one of the largest openly crowdsourced genome databases. In addition to the genetic data, members of *openSNP* annotate their data with additional trait data. There is no integration of further data sources into *openSNP*.

Despite the differences between *openSNP* and *Open Humans*, there is overlap of members that use both platforms, with *openSNP* members having additional non-genetic public data sets in *Open Humans*. By linking the public data sets across both platforms, both ecosystems can be enriched and members can avoid having to upload their data twice.

The connection of accounts is performed by each platform providing links to the same member on the other platform: The *openSNP* project for *Open Humans* asks members for permission to read their *Open Humans* username during the authentication phase. By recording a members *Open Humans* username, it becomes possible to link the public data sets on *Open Humans* to a given *openSNP member*. Furthermore, *openSNP* deposits a link to the public *openSNP* data sets in their *Open Humans* member account. So far over 250 people have taken advantage of linking their *openSNP* and *Open Humans* accounts to each other.

#### Genetic Data Augmentation

Most DTC genetic testing companies genotype customers using single-nucleotide polymorphism (SNP) genotyping technology, which genotypes a fraction of the total available sites in a human genome. As any two human genomes are more than 99 percent identical, these genotyped sites are carefully selected to capture human variation across global sub-populations. These sites (or genetic *variants*) can inform customers about their genetic ancestry, predict traits such as eye color, and determine susceptibility to some recessive diseases. While DTC testing may only genotype a tiny fraction of total sites available in the genome, it’s offered at a fraction of the price when compared to more comprehensive genotyping methods such as exome or genome sequencing. Until recently, individuals who wanted to know their genotypes at sites not covered by DTC testing needed to purchase a significantly more expensive genotyping test.

Genome-wide genotype imputation is an increasingly popular technique that offers a no-or low-cost alternative to comprehensive genotyping methods. In short, imputation is performed by scanning the entire genome in large intervals and using high-quality genotype calls from a large reference population to statistically determine a sample’s (or samples’) genotype likelihoods at missing sites based on shared genotypes with the reference population. Traditionally, genotype imputation has not been readily accessible to DTC customers because it entails a complex multi-step process requiring technical expertise and computing resources. Recently, the Michigan Imputation Server launched a free to use imputation pipeline [76]. The server was designed to be user-friendly and greatly lowered the barrier to entry for everyday DTC customers to have access to imputed genotypes.

As part of the Open Humans platform, *Imputer* is a participant-created project that performs genome-wide genotype imputation on one of a *Member’s* connected genetic data sources, such as *23andMe* or *AncestryDNA*. Firstly, *Imputer* must be authorized by a *Member*; once connected, the *Imputer* interface (https://openimpute.com) allows the *Member* to select which genetic data source they would like to impute and launches the imputation pipeline in one click. *Imputer* submits the imputation job to a queue on a server where the imputation is performed. Once the job has finished, the imputed genotypes are uploaded as a .*vcf* file and an email is sent to the *Member* notifying them that their data is available. *Imputer* makes it easy for *Members* to augment their existing genetic data sources using techniques that were previously dificult to access. The *Imputer* imputation pipeline was built using genipe [77] and uses the 1000 Genomes Project [78] genotype data as the reference population.

#### Re-use of Public Data for Understanding Health Behavior

A research team at the Universities of Copenhagen and Geneva, the Quality of Life (QoL) Technologies Lab, has been able to perform preliminary research using public data in Open Humans. As physical inactivity is one of the strongest risk factors for preventable chronic conditions [79], the QoL Lab’s goal is to leverage self-quantification data to assess and subsequently enhance the well-being of individuals and possibly, in the longterm, reduce the prevalence of some chronic diseases. At this stage, the QoL Lab has used the *Open Humans* public datasets of *Fitbit* and *Apple HealthKit* projects.

In *Open Humans*, individuals who donate public data uploaded from Fitbit and Apple HealthKit projects share with others the daily summaries taken with their *Fitbit* and *Apple* devices such as steps, resting heart rate (HR) and minutes asleep. The public datasets contain time series data from at least 30 members, each of whom decides whether to provide access to the aforementioned measurements. The number of records for each variable available in the *Open Humans* database varies since not all the devices record the same variables and participants may choose not to share a particular measurement (see Fig. 2).

**Figure 2.**
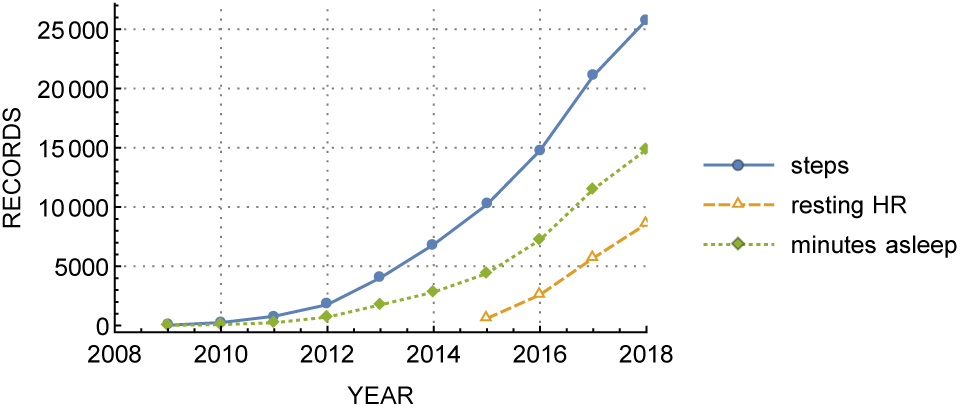
Self-quantification data from *Fitbit* project. Number of public records from January 2009 until October 2018 (cumulative total).

The QoL Technologies Lab team reports that access to public data has facilitated its research planning. While the number of public datasets is smaller in terms of the number of members who give this kind of access, they are very useful for running observational studies over long periods of time and can be used to prepare data cleaning and processing methods, which can then be applied to follow-up studies. As running studies on Open Humans and accessing private data as part of a research institution requires approval from an Institutional Review Board - a potentially lengthy process - the availability of the public data allows development and testing of methods during an earlier stage of the research process. A study is now being developed based on this preliminary work. Additionally, the QoL Lab has been granted ethics approval from University of Copenhagen in November 2018 (#504-0034/18-5000).

#### Data re-use in genetic data visualization research

With the increasing number of individuals engaging with their genetic data, including via direct-to-consumer products, there is a need for research into how individuals interact with this data to explore and understand it. The *Human-Computer Interaction for Personal Genomics* (PGHCI) project at Wellesley College and New York University has focused on exploring these questions. Research was initially conducted by creating visualizations of genetic data interpretations based on public genetic data sets and associated reports. The research initially recruited participants via Amazon Mechanical Turk to evaluate a set of visualizations; this approach, however, was not based on participants’ *own* information, which is preferred to improve experimental validity.

Open Humans provided an opportunity to work with individuals and their data in a manner that leveraged pre-existing genetic data for re-use in new research while minimizing privacy risks. A project, *GenomiX Genome Exploration*, was created in Open Humans that invited *Members* who had publicly shared their genetic data in Open Humans to engage with a custom visualization derived from their existing public data and associated interpretations. The study found various design implications in genome data engagement, including the value of affording users the flexibility to examine the same report using multiple views [80].

#### Personal Data Exploration

*Open Humans* aggregates data from multiple sources connected to individual *Members*. This makes it a natural starting point for a *Member* to explore their personal data. To facilitate this, *Open Humans* includes the *Personal Data Notebooks* project.

Through a *JupyterHub* setup (https://jupyterhub.readthedocs.io) that authenticates *Members* through their *Open Humans* accounts, *Members* can write *Jupyter Notebooks* [81] that get full access to their personal data in their web browser. This allows *Members* to explore and analyze their own data without the need to download or install specialized analysis software on their own computers. Furthermore, it allows *Members* to easily analyze data across the various data sources, for example combining data about their social media usage as well as activity tracking data from wearable devices. This allows *Members* to explore potential correlations such as whether a decrease in physical activity correlates with more time spent on social media.

As the notebooks themselves do not store any of the personal data, but rather the generic methods to access the data, they can be easily shared between *Open Humans Members* without leaking a *Member’s* personal data. This property facilitates not only the sharing of analysis methods, but also reproducible *n=1* experiments in the spirit of self-quantification.

To make these notebooks not only interoperable and reusable, but also findable and accessible [82], the sister project to the *Personal Data Notebooks* - the *Personal Data Exploratory* - was started. *Members* can upload notebooks right from their Personal Data Notebook interface to *Open Humans* and can publish them on the *Personal Data Exploratory* site with just a few clicks. The *Exploratory* publicly displays the published notebooks to the wider community and categorizes them according to the data sources used, tags and its content.

The categorization allows other *Members* to easily discover notebooks of interest. Notebooks written by other *Members* can be launched and run on a *Member’s* own personal data through the *Personal Data Notebooks*, requiring only a single click of a button. Through the close interplay between the *Personal Data Notebooks* and the shared notebook library of the *Personal Data Exploratory, Open Humans* offers an integrated personal data analysis environment that allows personal data to be disseminated in a private and secure way, while simultaneously growing a library of data exploration tools that can be reused by other *Members*, as shown in Figure 3.

**Figure 3.**
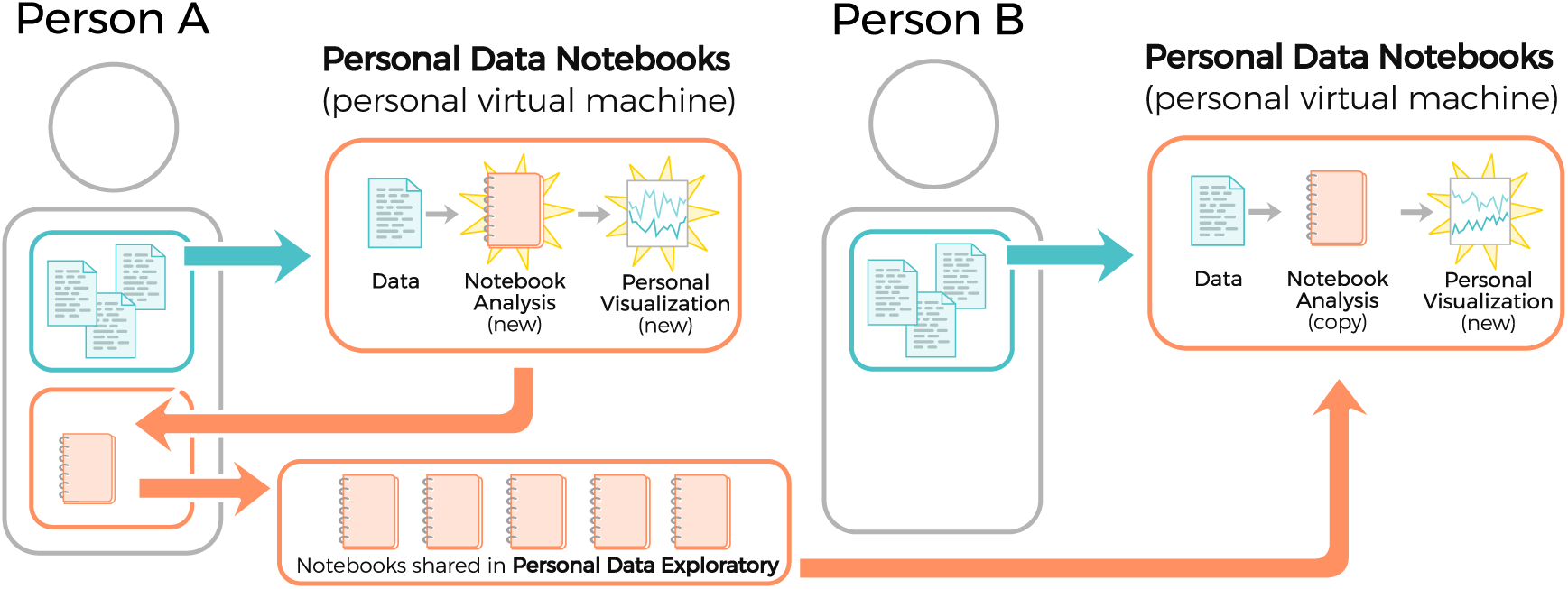
Personal Data Notebooks in Open Humans. Any *Member* (e.g. “Person A”, left), can create a *Notebook* to explore their personal *Data* using the *Personal Data Notebooks* project. They can then choose to share a *Notebook* via the *Personal Data Exploratory*. This allows another *Member* (e.g. “Person B”, right) to load a copy of the *Notebook* and run it, privately, to produce their own analysis.

#### Google search history analyzer and community review

The *Google Search History Analyzer* is a project that highlights the *Open Humans* community review process for *Projects*, demonstrating how this process can help improve not only a project that is reviewed, but also the infrastructure of *Open Humans*. The *Google Search History Analyzer* invites individuals to upload their Google Search History data, and analyze them in a quantitative way, through *Personal Data Notebooks*. Examples of analyses that users can do through the *Personal Data Notebooks* include retrieving graphs of their most common search terms and their daily or weekly evolution, as well as visualizing connections among their top search terms and their co-occurrence. One goal of this project is to raise awareness on the breadth and deeply personalized content that web searches might carry. Another long-term goal is to provide social scientists who are currently using web search history data for predicting social trends, e.g. unemployment [83], or interest in medical conditions [84], with the means to have access to a pool of individuals who can provide informed consent to the use of their search history data along with additional meta-data (e.g. demographic information, or survey questions), that could render their Google search history terms more informative.

*Open Humans* requires *Projects* have an “approval” to become visible and broadly available to *Members*. Prior to *Google Search History Analyzer*, this approval process was informal and internal; however, the sensitive data handled by this *Project* raised concerns regarding a need for a more formal and transparent review process – as web search history terms might carry highly personalized information, like personal interests, medical history, places a person visits, or even predictors for severe psychiatric conditions [85]. As a result, a community review process was developed for *Project* approvals going forward: new projects are shared with the larger community for public comments, inviting feedback from all *Members*. Project owners can reply to the feedback and improve their project accordingly, as well as seek help from other *Open Humans Members*. The community review continues until concerns, if any, have been resolved; no formal timeline for finding consensus exists, instead the process is adaptive to the levels of concern raised by the community members. If and when project approval occurs, this status is implemented by the administrators of the Open Humans platform. Project approval status can be re-considered at any time by opening a new review process, which may be done by any community *Member*.

As a result of this process, the *Google Search History analyzer* project was improved with added, documentation, increased clarity, and additional security implementations on the project side. Furthermore, it led to the implementation of a new feature on the *Open Humans* platform itself, enabling a project-specific override to prevent public data sharing by *Members* for this data – as requested by community review – thereby reducing the risk that these sensitive datasets might be publicly released by the *Members* accidentally.

## Discussion

Participatory/community science (also known as citizen science) is a growing field that engages people in the scientific process. But while participatory science keeps growing quickly in the environmental sciences and astronomy, its development in the humanities, social sciences, and in medical research lags behind [86], despite expectations that it will make inroads into those fields [60, 87]. Both barriers in accessing personal data that is stored in commercial entities as well as legitimate ethical concerns that surround the use of personal data contribute to this slower adoption in realms that rely on access to personal information [34, 36]. *Open Humans* was designed to address many of these issues—we discuss some of them in subsequent sections.

### Granular and Specific Consent

One often suggested way to mitigate the ethical concerns around the sharing of personal data in a research framework is by giving participants granular consent options [37]. In a medical context, most patients prefer to have granular control over which medical data to share and for which purposes [88, 89], especially in the context of electronic medical records [90]. Furthermore, the GDPR requires that organizations handling personal data give the individual granular consent options for how their data is used [91].

*Open Humans* strongly limits the platforms use of member data to an opt-in model, implementing a form of granular consent for data sharing and data use through the use of projects that *Members* can opt into. On a technical level, project organizers need to select the data sources they would like to access, and *Members* can give specific consent for that project’s activities. From the perspective of *Open Humans* this produces a format for granular consent regarding the data it manages, as each potential use of data in the platform is mediated by a specific project.

Additionally, projects on *Open Humans* need to adhere to the community guidelines. In addition to mandating clarity and specificity in consent, these guidelines require projects to inform prospective participants about the level of data access they would request, how the data would be used, and what privacy and security precautions they have in place. Authorization may be withdrawn at any time, at which point projects may no longer access de-authorized data. Furthermore, projects may receive notification of erasure requests made by participants who withdraw, should they opt to support these.

### Data portability

Much of health data is still stored in data silos managed by national institutions, sometimes further sub-categorized by diseases [92]. On an individual level, the situation is not much better: While medical data is usually stored in electronic records, much of a person’s data is now held by the companies that run social media platforms, develop smartphone apps, or purvey wearable devices [93]. This fragmentation—especially when coupled with a lack of data export methods—prevents individuals from authorizing new uses of their data.

Personal information management systems (PIMS) could help individuals in re-collecting and integrating their personal data from different sources [94]. The right to data portability, as encapsulated in Article 20 of the GDPR, has the potential to boost the adoption of such systems, as it guarantees individuals in the European Union a right to export the personal data they have provided to data holders in electronic and other useful formats. While Article 20 does not cover derived data, such as genetic information generated from biological samples [95], other personal data that is provided directly and thus subject to Article 20 can be highly valuable for individuals and research purposes. Additionally, Article 15 of the GDPR provides individuals with further rights to access and copies to such derived data, though without specific provisions for the format of such data. Both traditional medical research [96] as well as citizen science [97] have the potential to benefit from these data. By design, *Open Humans* works similar to a PIMS, as it allows individuals to bundle and collect their personal data from external sources. Like other PIMS, *Open Humans* is likely to benefit from any increase in data export, e.g. due to the GDPR.

While the availability of data export functions is a necessary condition for making PIMS work, it alone is not sufficient. PIMS need to support the data import on their end, either by supporting the file types or by offering support for the APIs of the external services. As file formats and APIs are not static, but can change over time, especially among popular services [98], a significant amount of effort is needed to keep data import functions into PIMS up to date. This cost keeps accumulating as the number of supported data imports keeps increasing. The modular, project-based nature of *Open Humans* allows the distribution of the workload of keeping integrations up to date, as data importers can be provided by any third party. Existing data imports on *Open Humans* already demonstrate this capability: Both the *Nightscout* and the *Apple HealthKit* data importers are examples of this. In the case of *Nightscout*, members of the diabetes community themselves built and maintain the data import into *Open Humans* to power their own data commons that overlays the *Open Humans* data storage. And the *HealthKit* import application was written by an individual *Open Humans Member* who wanted to add support for adding their own data.

### Enabling individual-centric research and citizen science

Open Humans provides several benefits for citizen science efforts and individual researchers who do not work in academia. The *OpenAPS* and *Nightscout Data Commons* highlighted in the results are prime examples of how Open Humans can enable such participant-led research.

To enable research done by non-traditional researchers, the project creation workflow of Open Humans includes information for project leaders about informed consent and other key considerations. It encourages project administrators to be clear about both data management and security in a thorough community guide https://www.openhumans.org/community-guidelines/#project. This guide includes best practice guidelines for data security as well as details on how to communicate to participants which data access is being requested and why. It emphasizes plain language and clarity.

To further the community’s sense of ownership in the *Open Humans* platform, participants are involved in the governance of the ecosystem. On a high level the community gets to elect a third of the members of the Open Humans Foundation board of directors, enabling them to exert direct influence on the larger direction of the platform.

Furthermore, *Members* of Open Humans are invited to participate in the approval of new projects that want to be shared on the platform via a community review process, as illustrated by the *Google Search History* project use case described above. This community review process parallels efforts made elsewhere to pursue participant-centred alternatives to institutional review boards [99], which at present include extremely limited input from community members. Indeed, traditional policies for project approval from an ethical standpoint have been repeatedly questioned [100], and even more so for the case of participant-centric research [101], due to inconsistent levels of engagement from non-academic members [102] and lack of participant protection and autonomy [101]. Notably the review process as implemented on *Open Humans* is less structured than traditional approaches, as it is performed by community members who choose to participate; self-selection for engagement may help maximize efficiency in a heterogeneous ecosystem. We hope this alternative design helps inform other projects seeking increased participant input in project review and oversight.

### Summary

*Open Humans* is an active online platform for personal data aggregation and data sharing that enables citizen science and traditional academic science alike. By leaving data-sharing decisions to individual members, the platform offers a way of doing personal data-based research in an iterative, ethically sensitive way and enables individuals to engage in science as both investigators and participants.

## Methods

The primary Open Humans web application, as well as data source *Projects* maintained directly by *Open Humans*, are written in Python 3 using the Django web framework. API endpoints, JSON and HTML data serialization, and OAuth2 authorization are managed by the *Django REST Framework* and *Django OAuth Toolkit* libraries. Web apps are deployed on *Heroku* and use *Amazon S3* for file storage. The *Personal Data Notebooks* JupyterHub project is deployed via *Google Cloud Platform*.

Two Python packages have been developed and distributed in the *Python Package Index* to facilitate interactions with our API: (1) *open-humans-api* provides Python functions for API endpoints, as well as command line tools for performing many standard API operations, (2) *django-open-humans* provides a reusable Django module for using *Open Humans* OAuth2 and API features.

Open Humans complies with GDPR and provides a live records of processing activities report at: https://www.openhumans.org/data-processing-activities/

## Availability of source code and requirements

- Project name: Open Humans
- Project home page: http://www.openhumans.org
- Operating system(s): Platform independent
- Programming language: Python3
- Other requirements: full list on GitHub https://github.com/openhumans/open-humans/
- License: MIT
- Project name: Open Humans API
- Project home page: https://open-humans-api.readthedocs.io/en/latest/
- Operating system(s): Platform independent
- Programming language: Python3
- Other requirements: full list on GitHub https://github.com/openhumans/open-humans-api
- License: MIT
- Project name: Django Open Humans
- Project home page: https://github.com/OpenHumans/django-open-humans
- Operating system(s): Platform independent
- Programming language: Python3
- Other requirements: full list on GitHub
- License: MIT

## Declarations

### List of abbreviations

API: application programming interface
CGM: Continuous Glucose Monitor
DTC: Direct to Consumer GDPR General Data Protection Regulation
IRB: Institutional Review Board
PIMS: Personal information management systems
QoL: Quality of Life

### Ethical Approval

Not applicable

### Consent for publication

Not applicable

## Author’s Contributions

BGT: Conceptualization, Data curation, Investigation, Methodology, Project administration, Software, Supervision, Writing - original draft, Writing - review & editing MA: Supervision, Writing - review & editing KA: Data curation, Software, Validation, Writing - original draft, Writing - review & editing MD: Software, Writing – review & editing VE: Data curation, Formal analysis, Investigation, Validation, Visualization, Writing - original draft, Writing - review & editing BG: Data curation, Resources, Software, Validation TH: Methodology, Resources, Software DL: Data curation, Formal analysis, Validation, Writing - original draft, Writing - review & editing OS: Investigation, Validation, Writing - review & editing ON: Investigation, Validation AT: Data curation, Software, Validation, Writing - original draft, Writing - review & editing JB: Conceptualization, Funding acquisition, Resources, Investigation, Project administration, Supervision MPB: Conceptualization, Data curation, Funding acquisition, Investigation, Methodology, Project administration, Resources, Software, Supervision, Writing - original draft, Writing - review & editing

## Acknowledgements

The authors would like to thank all members of the Open Humans community for their diverse contributions to Open Humans: Developing the process as well as platforms that link to Open Humans, sharing their personal data, advancing public knowledge sources, being active community members.

In this spirit, this manuscript was written as a community project done by and with Open Humans members following an open call for contributions.

In particular, the authors would like to thank Rosy Gupta, Manaswini Das, Jasmine Tamak and Tarannum Khan. They made valuable contributions as summer interns with Open Humans through the Outreachy internship program. The authors are grateful to Mike Escalante, who contributed in software development as well as mentoring for Outreachy.

The authors also would like to thank the reviewers - their input significantly improved the manuscript

